# CD45 limits Natural Killer cell development from common lymphoid progenitors

**DOI:** 10.1101/2023.04.17.537109

**Authors:** Lizeth G Meza Guzman, Craig D Hyland, Grace M Bidgood, Evelyn Leong, Zihan Shen, Wilford Goh, Tobias Kratina, Jai Rautela, James E Vince, Sandra E Nicholson, Nicholas D Huntington

## Abstract

The clinical development of Natural Killer (NK) cell-mediated immunotherapy marks a milestone in the development of new cancer therapies and has gained traction due to the intrinsic ability of the NK cell to target and kill tumour cells. To fully harness the tumour killing ability of NK cells, we need to improve NK cell persistence and overcome suppression of NK cell activation in the tumour microenvironment. The trans-membrane, protein tyrosine phosphatase CD45, regulates NK cell homeostasis, with genetic loss of CD45 in mice resulting in increased numbers of mature NK cells [1–3]. This suggests that CD45-deficient NK cells might display enhanced persistence following adoptive transfer. However, here we demonstrated that adoptive transfer of CD45-deficiency did not enhance NK cell persistence in mice, and instead, the homeostatic disturbance of NK cells in CD45-deficient mice stemmed from a developmental defect in the common lymphoid progenitor population. The enhanced maturation within the CD45-deficient NK cell compartment was intrinsic to the NK cell lineage, and independent of the developmental defect. CD45 is not a conventional immune checkpoint candidate, as systemic loss is detrimental to T and B cell development [4–6], compromising the adaptive immune system. Nonetheless, this study suggests that inhibition of CD45 in progenitor or stem cell populations may improve the yield of *in vitro* generated NK cells for adoptive therapy.

## Background

Leucocyte common antigen CD45 (Ly5; encoded by the *Ptprc* gene) is a prototypic transmembrane receptor-like protein tyrosine phosphatase (PTP) expressed on all nucleated hematopoietic cells and is required for normal lymphocyte development [7–11]. There are two proposed ligands for CD45, CD22 a B cell restricted transmembrane glycoprotein and galectin-1 a carbohydrate-binding protein with an affinity for β-galactosides [12–14]. To date, galectin-1 is the only physiological ligand that results in inhibition of PTPase activity upon CD45 engagement [13, 15]. Nonetheless, efforts to discover a ligand for CD45 that positively modulates PTPase activity are still ongoing.

CD45 has the ability to influence multiple signalling pathways through its various biochemical functions, the dominant one being dephosphorylation of the C-terminal negative regulatory tyrosine in the Src-family kinases, consequently activating Src kinases upon T and B cell receptor (TCR; BCR) engagement [4–6, 16]. It is therefore an indispensable positive regulator of T cell antigen receptor signalling in cell lines, mouse models and humans [17–20]. CD45 deficiency in humans’ results in a form of severe combined immunodeficiency (SCID) that is characterized by decreased T cell numbers, normal, decreased, or increased B cell numbers, and normal or increased numbers of NK cells [21–24].

In addition to its role in TCR and BCR activation [4–6, 16], CD45 has been reported to activate Syk, JNK and p38, downstream of Ly49D activation in NK cells [2, 25]. Lastly, CD45 has been shown to suppress Janus kinase (JAK) activity, resulting in negative regulation of cytokine receptor signalling [26].

In mice, the *Ptprc* gene consists of 34 exons that encodes an extracellular domain (exons 1-12), interdomain (exons 13-14), transmembrane domain (exon 15), and an intracellular cytoplasmic domain (exons 16-34). T, B and NK cells express multiple CD45 isoforms, that are differentially expressed during developmental stages [10]. CD45 isoform switching occurs as lymphoid cells differentiate into T, B and NK cells, with less mature populations expressing larger isoforms, suggesting a lower activation threshold compared to more mature populations expressing the smaller isoforms [27].

Three CD45-deficient models have been generated in mice by independently targeting exons 6 [4], 9 [5] and 12 [6]. All three models display defects in thymic development mediated by increased apoptosis and dysfunctional pre-TCR and TCR signalling [16]. Deletion of exon 6 resulted in complete abrogation of CD45 expression on all B cells, while a small population of thymic T cells (3-5%) retained CD45 expression [4]. Deletion of exon 9 resulted in complete loss of CD45 on B and T cells [5]. Lastly, targeting exon 12, which encodes part of the extracellular domain in all isoforms, resulted in complete loss of CD45 surface expression on all cell types [6]. Although this CD45 null mouse strain (*Cd45^−/−^*) lacks membrane bound CD45, immunoblotting revealed the presence of a truncated protein (∼150 kDa) [6]. Despite this, the exon 12-targeted *Cd45^−/−^* phenotype is identical to the exon 9-targeted model.

CD45-deficient mice lack mature T and B cells [4–6, 16]. T cell loss is due to dysfunctional TCR signalling leading to a reduction in double positive and single positive thymocytes, as well as a defect in negative selection following antigen stimulation [28]. Thus, the CD45-deficient T cells that do develop and reach the periphery are highly autoreactive. B cell development halts at T2 transitional stage in the spleen as B cells fail to express IgD [29].

Although CD45 is required for T and B cell development, this does not appear to be the case for NK cells [2, 30]. Splenic NK cells in *Cd45^−/−^* mice were increased 4-5-fold and actively dividing, compared to NK cells in control mice, suggesting that CD45 is a regulator of NK cell homeostasis, functioning to limit NK cell numbers [1, 2]. Additionally, splenic NK cells from *Cd45^−/−^* mice retained cytotoxic ability but exhibited defective IFNγ production [2, 3].

The *in vivo* expansion and/or persistence of NK cells in CD45-deficient mice suggests that CD45 is a potential target for immunotherapies designed to enhance NK cell anti-tumour activity. Adoptive NK cell therapies are considered safe, with no cytokine release syndrome or graft-vs-host disease observed in clinical trials so far and have been successfully used against relapsed or refractory CD19-positive cancers (non-Hodgkin’s lymphoma or chronic lymphocytic leukemia [31, 32]. Targeting CD45 in NK cell-based immune cell therapy has the potential to bypass the severe T and B cell immunosuppression expected with whole body inhibition of CD45 activity. Here we investigated the role CD45 plays in regulating NK cell homeostasis and explored the potential of inhibiting CD45 activity for NK cell immunotherapy.

## Methods

### Mice

All animal experiments followed the National Health and Medical Research Council (NHMRC) Code of Practice for the Care and Use of Animals for Scientific Purposes guidelines and were conducted in accordance with the regulatory standards approved by the Walter & Eliza Hall Animal Ethics Committee (AEC2019.034, AEC2018.040, AEC2021.011 and SABC) and Monash University Animal Ethics Committee (25004, 22111). All strains (**Table 1**) were maintained on a C57BL/6 background and bred at either the Walter & Eliza Hall Institute or Monash University.

**Table 1.** Genetically modified mouse strains.

| Strain<sup>1</sup> | AKA<sup>2</sup> | Description | Reference<sup>3</sup> |
| --- | --- | --- | --- |
| Cd45tm1Tyb | *Cd45*<sup>-/-</sup> | Disruption of Cd45 gene by insertion of a neomycin gene | [6] |
| Mcl-1<sup>fl/fl</sup> Ncr1-Cre | *Mcl1*<sup>fl/fl</sup> *Ncr1*<sup>iCre/+</sup> | NK lymphopenic mice | [33] |
| IL-15(-/-) | IL-15<sup>-/-</sup> | C57BL/6 mice genetically deficient in interleukin 15 | [34] |
| IL-15tg | IL-15 Tg<sup>T/+</sup> | IL-15 transgene where three posttranscriptional checkpoints were eliminated. Transgene overexpression achieved by the MHC class I promoter | [35] |
| B6 Cd45.1 | Ly5.1/J | C57BL/6 congenic strain carrying the Ptpca pan leukocyte marker commonly known as CD45.1 or Ly5.1 | [36-38] |
| Ly5.1Jx C57BL/6 | Ly5.1x5.2 | Cross between B6.SJL-Ptpca<a>Pepc<b>/BoyJ (Ly5.1) acquired in 2017 to C57BL/6J | [37] |
| C57BL/6J | C57BL/6 | Acquired from JAX in 1989. | [39] |
| Rag2<sup>-/-</sup> common gamma chain<sup>-/-</sup> | *Rag2*<sup>-/-</sup> γc<sup>-/-</sup> | Completely alymphoid (T-, B-, NK-) mice | [40] |
| B6.129S7-Rag1<tm1Mom> | *Rag1* | Mice homozygous for the Rag1tm1Mom mutation produce no mature T cells or B cells | [41] |
| Ptpcr<sup>fl/fl</sup> | *Cd45*<sup>fl/fl</sup> | Generated by Monash Genome Modification Platform | This publication |
| Nkp46<sup>iCre/wt</sup> | *Ncr1*<sup>iCre/+</sup> | “Knock-in” mice containing the iCre recombinase gene under control of the Ncr1 gene promoter | [42] |
| Ptpcr<sup>fl/fl</sup> Ncr1<sup>Cre/+</sup> | *Cd45*<sup>fl/fl</sup> *Ncr1*<sup>Ki/+</sup> | Mice with conditional deletion of CD45 in Nkp46 expressing cells | This publication |

<sup>1</sup>Strain name as appeared in first publication. <sup>2</sup>Common strain name. <sup>3</sup>Strain generation and characterization

### Generation of mice with conditional deletion of *Cd45* (*Ptprc*) in NK cells

Mice with a floxed Cd45 allele (*Cd45^fl/fl^*) were generated on a C57BL/6 background by the Monash Genome Modification Platform (**Supplementary Figure 1**). Clustered Regularly Interspaced Short Palindromic Repeats (CRISPR) technology was used to generate mice carrying *Cd45* conditional alleles (loxP). In short, CRISPR ribonucleoproteins (RNPs; Cas9 protein in complex with sgRNAs targeting sequences flanking *Cd45* exon 14) and a donor repair template (homologous single-stranded oligonucleotide with mutations to insert loxP sites) were microinjected into fertilized one-cell stage embryos. sgRNA guide and donor template sequences are provided in **Supplementary Figure 1B & C**. Cre-lox deletion of exon 14 results in a premature stop codon in exon 15. Furthermore, translation of the compromised mRNA sequence predicts a truncated CD45 protein, which lacks a transmembrane and cytoplasmic domain and is expected to be non-functional (**Supplementary Figure 1D**). Established *Cd45^fl/fl^* mice were crossed to *Ncr1^iCre/+^* mice to conditionally delete *Cd45* from mature NK cells (*Cd45^fl/fl^ Ncr1^iCre/+^*). These strains were maintained on a C57BL/6 background and bred at Monash University animal facilities (Clayton).

### Organ processing

Blood, BM, liver, lungs, and spleen were collected from 6-8-week-old mice for further analysis. Each organ was processed to a single cell suspension as follows:

**Blood:** Retro-orbital, mandibular or cardiac bleeds were collected in Microvette® 500 K3 EDTA tubes (Sarstedt), transferred to a 5 mL polypropylene tube (Falcon), and adjusted to a total volume of 1 mL with PBS. The cell suspension was under-layered with 2 mL of Histopaque-1077 (Sigma Aldrich) and centrifuged at 300 *g* for 15 min at room temperature (RT). Leukocytes can be found at the interface layer, these were then transferred to a 10 mL falcon tube, washed twice with ice-cold phosphate-buffered saline (PBS; Gibco) and resuspended in FACS buffer [PBS, 2% FBS, 1 mM ethylenediaminetetraacetic acid (EDTA)]. Enriched leukocytes were used in subsequent experiments.

**Bone Marrow:** Unless otherwise specified, femurs and tibias were collected, and BM flushed into a 10 mL falcon tube using a PBS filled syringe. Cell suspensions were passed through a 70 µm cell strainer and centrifuged at 300 *g* for 5 min at 4°C. The cell pellet was resuspended in 1 mL of red cell removal buffer (RCRB; 156 mM NH_4_Cl, 11.9 mM, NaHCO_3_ and 0.097 mM, EDTA) and incubated for 5 min at RT. Cells were washed twice with ice-cold PBS and resuspended in FACS buffer. BM single cell suspensions were used in subsequent experiments.

**Livers** were forcefully passed through a 70 µm cell strainer with PBS, sieved tissue collected in a 50 mL falcon tube and centrifuged at 300 *g* for 5 min at 4°C. Pellets were resuspended in 10 mL of 33.75% v/v Percoll (made from 90% isotonic Percoll; Sigma Aldrich) and centrifuged at 300 *g* for 30 min at RT, with low or no break/acceleration. Cells were washed twice with ice-cold PBS, centrifuged at 300 *g* for 5 min at 4°C, resuspended in 5 mL RCRB, and incubated for 5 min at RT. Cells were washed twice with ice-cold PBS and resuspended in FACS buffer. Liver single cell suspensions were used in subsequent experiments.

**Lungs** were minced at RT into small fragments. Minced lungs were thoroughly mixed with 5 mL digestion buffer (1 mg/mL collagenase IV and 30 µg/mL DNase I in PBS) and incubated for 30 min at 37°C. To further dissociate cells, the digested tissue was forcefully passed through a 70 µm cell strainer with PBS, cell suspension was centrifuged at 300 *g* for 5 min at 4°C. Cell pellet was resuspended in 5 mL of RCRB, incubated for 5 min at RT and then washed twice with ice-cold PBS.

**Spleens** were forcefully passed through a 70 µm cell strainer with PBS, collected in a 10 mL falcon tube and centrifuged at 300 *g* for 5 min at 4°C. The cell pellet was resuspended in 1 mL of PBS and passed through a 70 µm cell strainer with PBS, adjusting the volume to 10 mL in a 10 mL falcon collection tube, prior to centrifugation at 300 *g* for 5 min at 4°C.

### Flow cytometry analysis

Cell proliferation and intracellular IFN**γ** data was collected on a FACSVerse (BD Biosciences) using BD FACSuite software. NK cell progenitor data was collected on a FACSymphony (BD Biosciences) using BD FACS Diva software. Data for all other experiments was collected on a BD LSR Fortessa X-20 using BD FACS Diva software. All analysis and statistics were performed using FlowJo v10 software and Prism GraphPad, respectively.

Flow cytometric analysis of viable cells was performed by excluding debris (FSC-A; Forward Scatter-Area low events), doublets (FSC-A vs FSC-H; Forward Scatter-Height) and dead cells (Dead-cell indicator dye negative). Cell subsets were gated based on surface marker expression (**Table 2**).

**Table 2.**
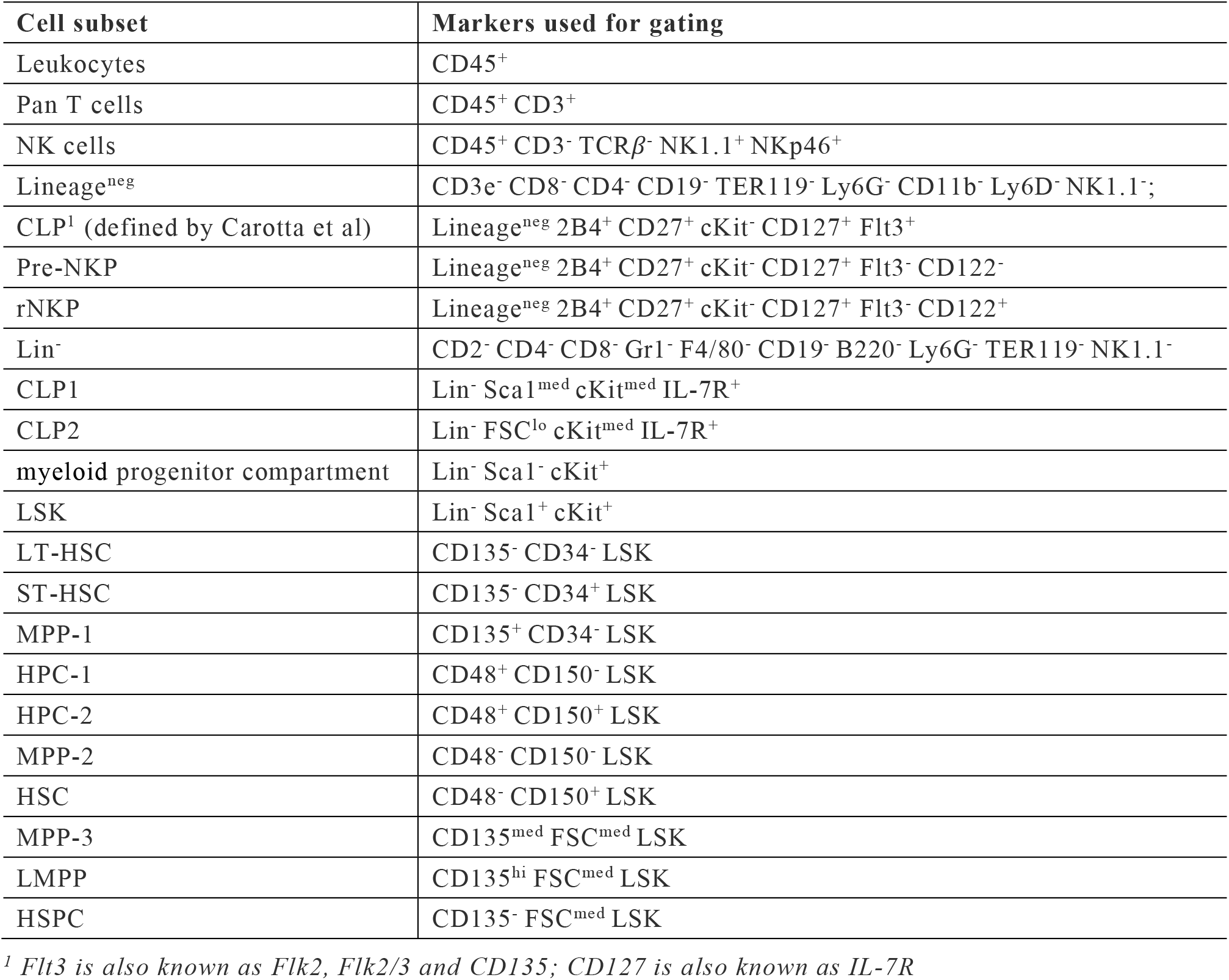
Markers used to gate various immune populations by flow cytometry.

### Production of IFNγ by splenic NK cells

Spleens were processed and intracellular IFN**γ** measured as previously described [43]. In short, splenocytes of 6-8-week-old mice were stimulated with 20 ng/mL recombinant mouse IL-12 (Peprotech) in combination with 200 ng/mL recombinant mouse IL-18 (Peprotech) or 50 ng/mL of recombinant human IL-15 (Miltenyi) for 4 h prior to fixation/permeabilization for detection of intracellular IFN**γ** by flow cytometry.

### NK cell purification

Splenic NK cells from 6- to 8-week-old adult mice were purified using the Miltenyi Biotec NK Cell Isolation Kit mouse as per the manufacturer’s instructions.

### *In vitro* proliferation assays

NK cell proliferation studies, including cohort number and mean division number determination, were performed as described [44], and based on previously published methods using T cells [45–47]. In short, purified NK cells are incubated with 0.1 nM of CellTrace Violet (CTV; Invitrogen™) in phosphate buffered saline (PBS) supplemented with 0.1% bovine serum albumin (BSA; Sigma Aldrich) for 20 min at 37°C. Labelled cells were subsequently washed twice with ice-cold NK complete media [IMDM (Gibco-BRL, Grand Island, NY) containing 10% v/v heat-inactivated FBS, 1% v/v penicillin-streptomycin, 1× GlutaMAX™ (Gibco-BRL, Grand Island, NY), 55 μM β-mercaptoethanol] to quench the labelling reaction. Labelled NK cells (4,000-10,000 per well) were seeded into 96-well round-bottom plates and cultured at 37 °C in 5% CO_2_. A mixture of propidium iodide (PI; 200 nM; Sigma-Aldrich) and 123count eBeadsTM (Invitrogen, 5005 beads/well) was added to cultures prior to flow cytometric analysis. Cells were analysed daily by flow cytometry.

### Generation of bone marrow (BM) chimeras

Bone marrow chimeras were generated by lethally irradiating (2 x 550 rads) host mice and reconstituting by I.V. injection into the tail vein with 6 x 10^6^ donor BM cells for 100% chimeras, or 3 x 10^6^ control donor BM and 3 x 10^6^ experimental donor BM for mixed chimeras. Generally, host and donor mice express allelic variants of the pan-hematopoietic cell marker CD45, also known as CD45.1 (Ly5.1) and CD45.2 (Ly5.2). Donor BM was collected from femur, tibia, pelvis, radius ulna, humerus, and cervical vertebrae by crushing the bones in a mortar with a pestle and PBS. BM suspension was passed through a 70 µm cell strainer, centrifuged at 300 g for 5 min at 4°C, resuspended in PBS and cells counted, prior to injection. Immediately post-irradiation, neomycin treated water (2 mg/mL neomycin sulphate in drinking water; Sigma) was made available for mice for up to 3 weeks. Reconstitution of the hematopoietic compartment was checked at 6-weeks post-bone marrow transplantation. Peripheral blood was obtained by retro-orbital bleed using a sterile hematocrit capillary tube (Sarstedt). Post-processing, blood samples were stained with anti-CD45.1 and anti-CD45.2 for flow cytometric analysis of the leukocyte compartment.

### Adoptive transfer of mouse NK cells

Splenic NK cells from 6-8-week-old mice were enriched from single cell suspensions by negative depletion. Splenocytes were labelled with biotinylated antibody cocktail (CD3, CD4, CD8, MHC-II, Ly6G, 84/80, TER119 AND CD19; antibody details in **Supplementary Table 1**) followed by enrichment using MagniSort^TM^ streptavidin negative selection beads (Invitrogen™). NK cells enriched from C57BL/6 (CD45.2) and *Cd45^−/−^*mice (∼70% purity) were labelled with 0.1 nM CTV. A portion of the cells were labelled with fluorescently conjugated antibodies against NK cell markers in combination with a mixture of PI (200 nM) and 123count eBeadsTM (5005 beads/well) prior to flow cytometric analysis. NK cells were counted, mixed (1:1) and injected into the tail vein of host mice.

## Results

Mice lacking CD45 have reduced numbers of mature T and B cells, but increased numbers of mature M2 stage NK cells with enhanced *in vitro* cytotoxic capability [1, 2]. However, previous studies did not address whether the increased NK cell maturation in *Cd45^−/−^* mice was related to IL-15 signalling, or at what stage in NK development CD45 limited expansion of the NK cell pool.

IL-15 is critical for NK cell survival and proliferation and contributes to IFNγ production [48–52]. Given that CD45 is also known to negatively regulate cytokine receptor signalling [26], we explored the interplay between IL-15 responses and the phenotype of CD45-deficient mice. Initially, splenic cells from wild-type C57BL/6 (CD45.2) and *Cd45^−/−^* mice were stimulated *ex vivo* with IL-12 and IL-18, or IL-15 cytokines. *Cd45^−/−^* NK cells produced slightly more IFNγ in response to IL-15 treatment, compared to wild-type NK cells. No differences were observed with IL-12 and IL-18 treatment. (**Figure 1A**). Degranulation, as measured by CD107a expression, was not different in IL-15 treated CD45^−/−^ NK cells (data not shown).

**Figure 1.**
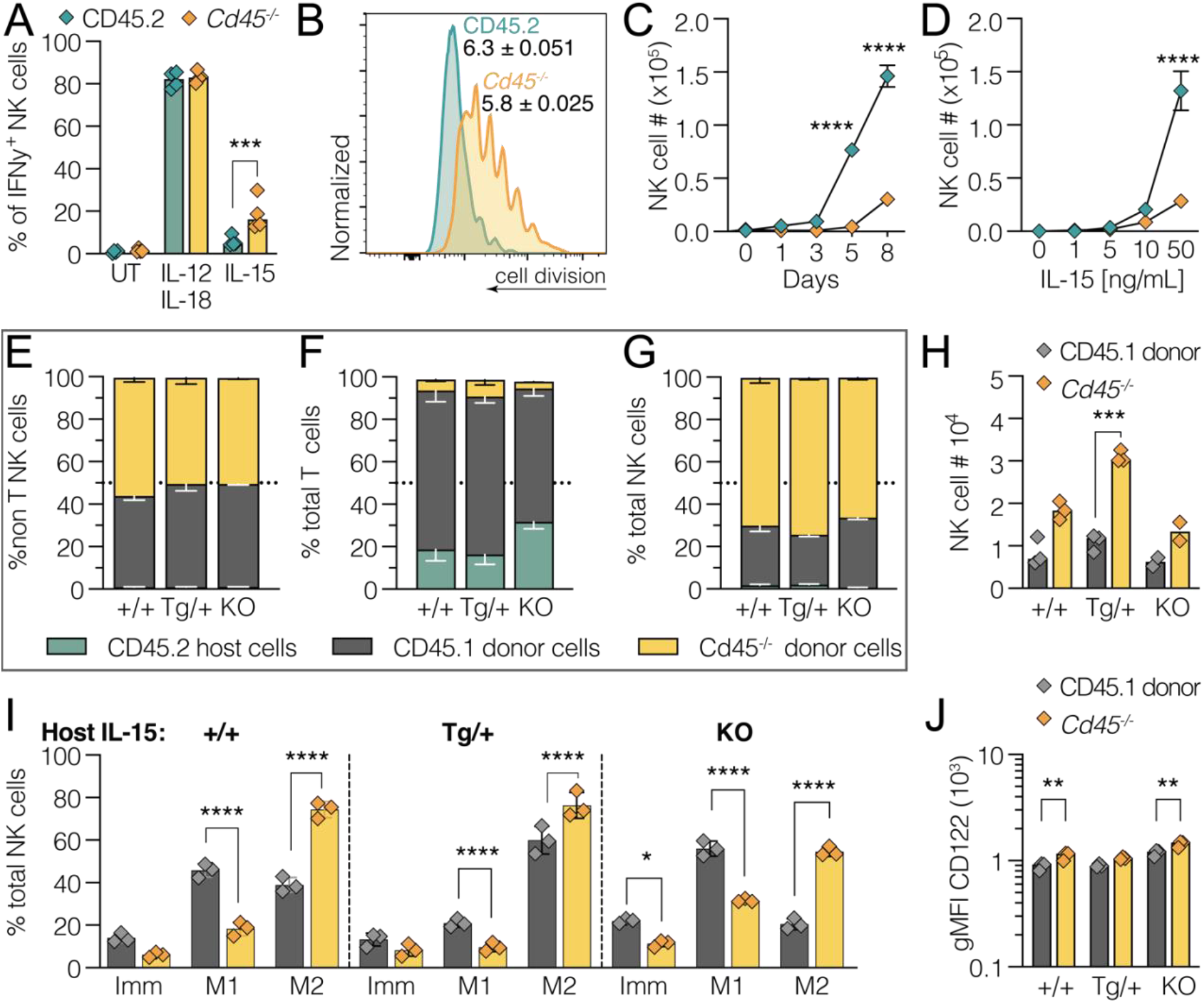
The in vivo expansion of CD45 ^−/−^ NK cells is not dependent on IL-15. **(A)** Splenocytes from either control C57BL/6 mice (CD45.2; green) or Cd45 ^−/−^ mice (yellow), were untreated (UT) or treated for 4 h with cytokines, and NK cells (CD3 ^−^TCRß^−^NK1.1^+^NKp46^+^) analysed for intracellular IFN**γ** production. Percentage of IFN**γ**^+^ NK cells. Representative data of two separate experiments. **(B - D)** Splenic NK cells from control C57BL/6 mice (CD45.2; green) or Cd45^−/−^ mice (yellow) were isolated and labelled with CellTraceViolet (CTV; 5 µM), prior to culturing for 8 days with recombinant human IL-15 (0, 1, 5, 10, 50 ng/mL). Viable cell numbers were analysed daily. **(B)** Example histograms depicting CTV-labelled NK cell division peaks following 8-days culture in 50 ng/mL IL-15. **(C)** Absolute numbers of NK cells with 1 to 8 days culture in 50 ng/mL hIL-15. **(D)** Absolute numbers of NK cells at day 8 with titration of IL-15 levels. Each point represents cells from an individual mouse. Total of 6 mice per genotype over 3 separate experiments. Note the data corresponding to 8-days culture in 50 ng/mL IL-15 is plotted in both C and D. **(E-J)** Bone marrow was harvested from control C57BL/6/CD45.1 (Ly5.1; grey) and Cd45 ^−/−^ (yellow) mice, mixed in a 1:1 ratio and a single cell suspension transplanted into lethally irradiated IL-15 Tg^+/+^, IL-15 Tg^T/+^ and IL-15^−/−^ CD45.2/Ly5.2 host mice to generate competitive bone marrow chimeras. 8-weeks post-transplantation, spleens were analysed for reconstitution by CD45.1 and Cd45 ^−/−^ donor cells: **(E)** % non-T/NK cells (CD3^−^NK1.1^−^), **(F)** % pan T cells (CD3^+^), (**G)** % NK cells (CD3^−^/NK1.1^+^/Nk46^+^), and **(H)** total NK cell numbers. (**I)** NK maturation profile (gating on CD11b and KLRG1), and (**J)** surface expression of CD122. Competitive chimeras were generated twice, with at least 5 hosts per genotype. E-G n= 5, H-J n=3 representative experiment. Significance was determined by two-way ANOVA with Sidak’s multiple comparisons test (**** <0.0001).

### *In vitro* proliferation of *Cd45^−/−^* NK cells is compromised in response to IL-15

To further assess the role of CD45 downstream of IL-15 signalling, splenic NK cells were purified from wild-type and *Cd45^−/−^* mice, labelled with CTV, and NK cell division tracked over 8 days in the presence of various hIL-15 concentrations (**Figure 1B - D**). Wild-type NK cells divided on average 6.3 times (50 ng/mL; 8 days), consistent with previous data [44]. In contrast, *Cd45^−/−^*NK cells were not able to reach division 7, and instead divided ∼5.8 times, with most cells achieving less divisions (**Figure 1B**). The normalized *Cd45^−/−^*NK cell cohort number (relative to initial number of cells that undergo division) was also decreased (0.226±0.043 vs 0.917±0.0367; *Cd45^−/−^* vs wild-type). At day 5 and day 8, the total numbers of *Cd45^−/−^*NK cells were significantly reduced compared to wild-type NK cells (∼4.6-fold in 50 ng/mL IL-15) (**Figure 1C & D**). This result was not consistent with the described *in vivo* expansion of the NK cell pool in *Cd45^−/−^*mice [2].

### CD45 regulation of NK cell homeostasis and maturation is not IL-15 independent

Although the ability of *Cd45^−/−^* NK cells to respond to IL-15 was compromised *ex vivo*, it was not clear whether homeostatic maintenance of NK cells *in vivo* (which also requires IL-15), would be impacted. To investigate this, BM cells from *Cd45^−/−^* (Ly5.1/2^−/−^) and CD45.1 (Ly5.1) mice were mixed (1:1) and transplanted into lethally irradiated wild-type mice (IL-15 Tg^+/+^ littermates), IL-15 transgenic mice (IL-15 Tg^T/+^), and IL-15 deficient mice (IL-15^−/−^) (all hosts: CD45.2/Ly5.2). Note, IL-15 deficiency is not complete in IL-15^−/−^ hosts, as transplanted cells can produce IL-15. Considering the lack of mature T cells and expansion of the NK cell compartment in *Cd45^−/−^* BM chimeras [4–6], reconstitution was assessed in the splenic CD3^−^NK1.1^−^ population (non-T/NK cells), which showed equal reconstitution by *Cd45^−/−^* and CD45.1 donor cells (**Figure 1E**).

As expected, less than 10% of reconstituted T cells (CD3^+^) were of *Cd45^−/−^* donor origin (**Figure 1F**). Lastly, the NK cell compartment in all recipients was dominated by *Cd45^−/−^* donor cells (>60%), with a corresponding increase in *Cd45^−/−^*NK cell number (**Figure 1G & H**). Both wild-type and *Cd45^−/−^* NK donor cell numbers were increased in IL-15 Tg^T/+^ host mice, but *Cd45^−/−^* NK donor cells were not selectively enhanced by IL-15 (1.2-fold for *Cd45^+/+^;* 1.6-fold for *Cd45^−/−^*) (**Figure 1H**). The reconstituted *Cd45^−/−^*NK cells skewed towards an M2 maturation profile in all hosts. This was independent of IL-15 levels (**Figure 1I**), and consistent with previous reports in chimeric wild-type mice [2]. Similarly, *Cd45^−/−^* cells displayed elevated levels of the IL-15 receptor beta subunit (IL-2Rβ; CD122), and this was also independent of IL-15 levels (**Figure 1J**).

In contrast to the reduced *in vitro* proliferation of *Cd45^−/−^*NK cells (**Figure 1B-D**), the enhanced *in vivo* expansion of the *Cd45^−/−^*NK cell compartment was recapitulated in the mixed bone marrow chimeras. Changes in IL-15 levels had no impact on either the expansion of the *Cd45^−/−^*NK cell compartment or the skewing towards mature M2 effector NK cells.

### Adoptively transferred *Cd45^−/−^* NK cells display reduced expansion in lymphocyte depleted mice

Although the competitive bone marrow chimeras confirmed that expansion of the NK cell compartment in *Cd45^−/−^* mice was intrinsic to hematopoietic cells, the chimeras did not address whether expansion was intrinsic to the NK cell lineage or resulted from a defect at an earlier stage of hematopoiesis. Competitive, adoptive NK cell transfers were performed to determine whether CD45-deficient NK cells were able to expand and accumulate *in vivo*. *Cd45^−/−^* and *Cd45*^+/+^ NK cells were purified, labelled with CTV, mixed (1:1) and adoptively transferred into NK cell-deficient mice (*Mcl1^fl/fl^ Ncr1^iCre/+^*), T and B cell-deficient mice (*Rag-1^−/−^*) or completely alymphoid mice (Rag-2^−/−^γc^−/−^).

In contrast to the bone marrow chimeras (**Figure 1G & H**), the expansion of adoptively transferred *Cd45^−/−^* NK cells was significantly impaired in *Rag-2^−/−^γc^−/−^* and *Rag-1^−/−^*hosts, with less *Cd45^−/−^*cells undergoing less cell divisions, than adoptively transferred CD45.1 NK cells (**Figure 2A-C**). This was consistent with a reduced contribution by transferred *Cd45^−/−^* cells to the NK cell compartment in spleen and lung, compared to CD45.1/Ly5.1 NK cells (for example, 33% vs 66% respectively, in *Rag-2^−/−^γc^−/−^*) (**Figure 2B-C**). To further examine the capacity of *Cd45^−/−^* NK cells to persist *in vivo*, NK cells were adoptively transferred into NK cell-deficient mice (*Mcl1^fl/fl^ Ncr1^iCre/+^*). Again, the relative contribution of *Cd45^−/−^* cells to the NK cell compartment was significantly less than observed for wild-type NK cells (**Figure 2D**).

**Figure 2.**
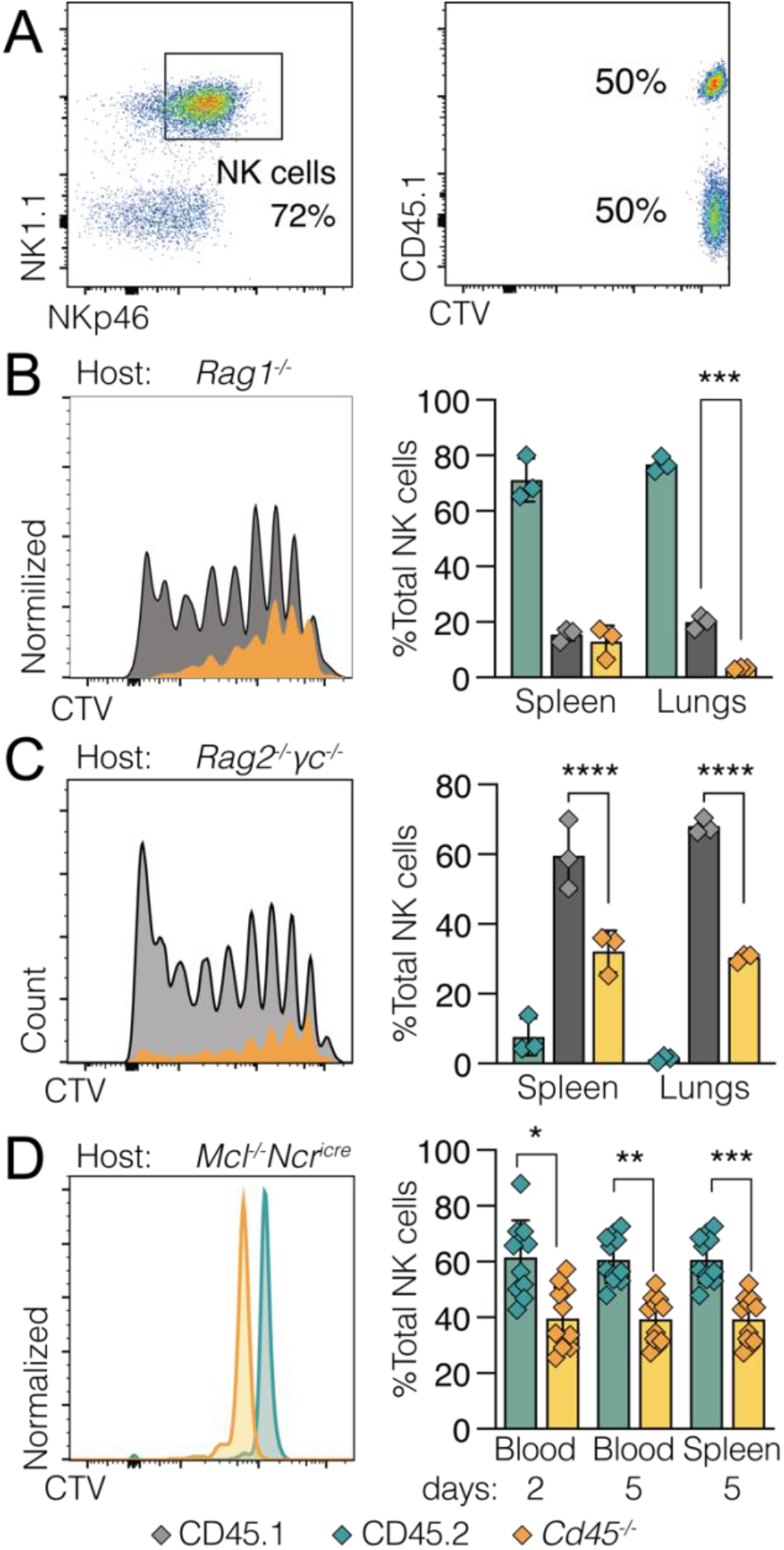
Loss of CD45 compromises the ability of NK cells to expand following adoptive transfer. **(A-C**) Splenic NK cells were enriched from CD45.1 (Ly5.1; grey) and Cd45 ^−/−^ (Ly5.1/2 null; yellow) mice by negative depletion, labelled with CTV, and mixed in a 1:1 ratio for I.V. injection into T and B cell-deficient (Rag-1^−/−^), or completely alymphoid (Rag-2^−/−^*γ*c^−/−^) host mice (CD45.2; green). **(A)** Confirmation of NK cell enrichment (left panel), and 1:1 ratio of CTV labelled cells prior to injection (right panel). **(B)** Rag-1^−/−^ and **(C)** Rag-2^−/−^*γ*c^−/−^ mice were sacrificed 6-days post-adoptive transfer, and spleens collected for analysis of NK cell proliferation (left panels) and relative proportions of donor and host cells in the NK cell compartment (right panels). Representative data (n=3) from 2 independent experiments. **(D)** Splenic NK cells were enriched from Cd45.2 (Ly5.2; green) and Cd45 ^−/−^ (Ly5.1/2 null; yellow) mice by negative depletion, labelled with CTV, and mixed in a 1:1 ratio for I.V. injection into NK cell-deficient mice (Mcl1^fl/fl^Ncr1^Ki/+^). 2 days post-adoptive cell transfer, blood samples were analysed for frequency of transferred NK cells. At day-5 post-adoptive cell transfer, blood and spleens were collected and analysed for frequency of transferred NK cells (right panel). Representative histograms are shown on the left. Data are combined from 3 independent experiments with each symbol representing an individual mouse. Statistical significance determined by two-way ANOVA with Sidak’s multiple comparisons test.

These observations were in sharp contrast to the original reported expansion of NK cells [2], and the bone marrow chimera data in **Figure 1**, which suggested that deletion of CD45 conferred an intrinsic proliferative advantage to NK cells. Instead, these data indicated that CD45 acted prior to NK cell development. To confirm this hypothesis, mice were generated with conditional deletion of *Cd45* in NK cells.

### Conditional deletion of CD45 in NK cells does not disrupt NK cell homeostasis, but does skew maturation towards mature M2 cells

CRISPR/Cas9 gene editing was used to generate mice carrying *Cd45/ptprc* conditional alleles (loxP). Mice carrying the floxed *Cd45* allele (*Cd45^fl/fl^*) were crossed to mice containing the cre recombinase gene under control of the *Ncr1* gene promoter (encoding NKp46; Ncr1^iCre/+^), to generate *Cd45^fl/fl^Ncr1^iCre/+^* mice. This resulted in deletion of C*d45* exon 14 from the immature stage of NK cell development (**Supplementary Figure 1**). Lack of CD45 surface expression on *Cd45^fl/fl^Ncr1^iCre/+^*NK cells was confirmed by flow cytometry (**Figure 3A**).

**Figure 3.**
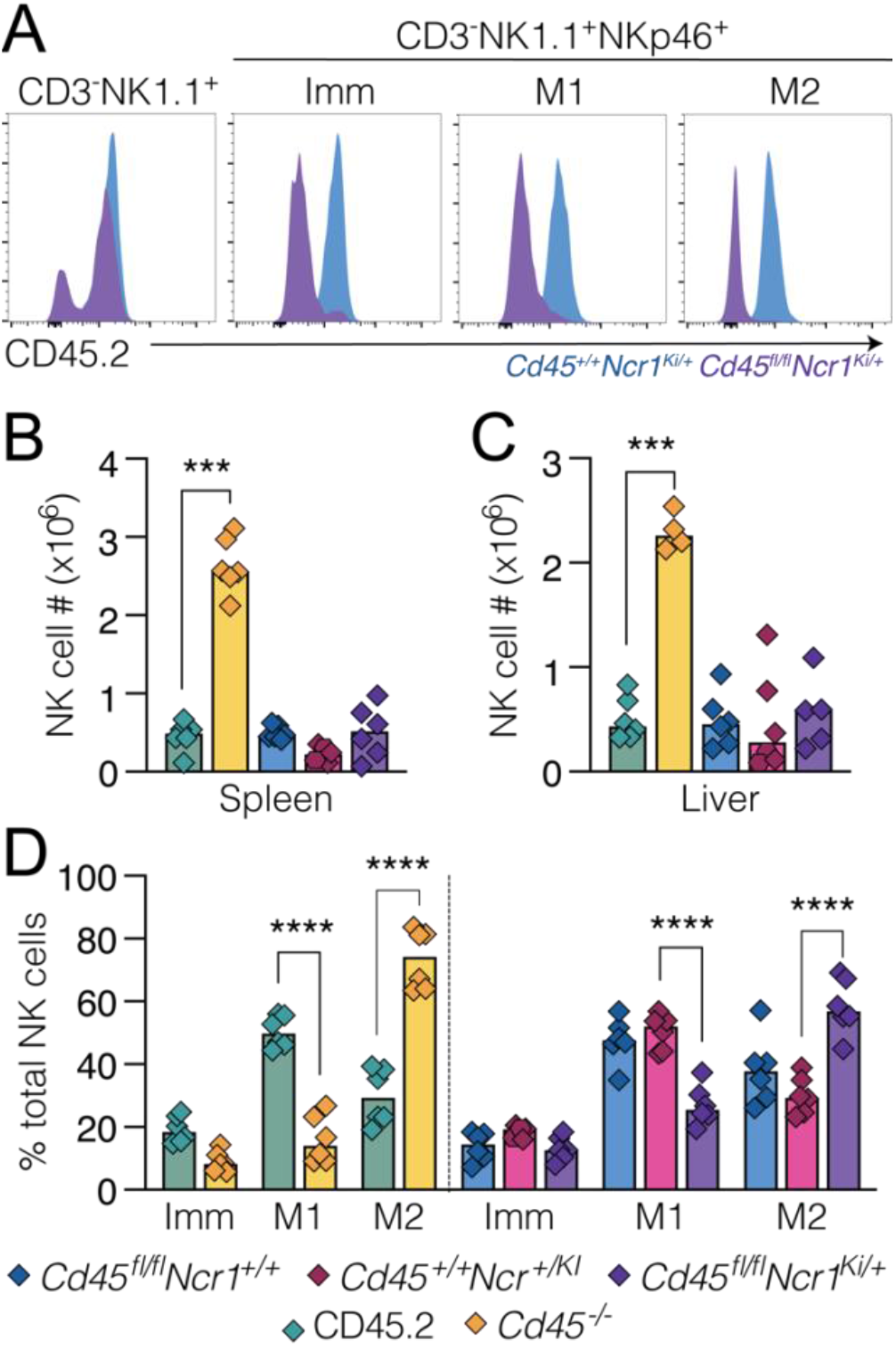
Loss of CD45 in NK cells disrupts NK cell maturation but not homeostasis. **(A)** Flow cytometric analysis confirming conditional deletion of CD45 in immature and mature NK cells. Bone marrow cells from control mice (Cd45^+/+^Ncr1^iCre/+^; blue) and mice with NK deletion of Cd45 (Cd45^fl/fl^Ncr1^iCre/+^; purple) were gated on NK precursor markers (CD3 ^−^NK1.1^+^), NK cell markers (CD3^−^ NK1.1^+^NKp46^+^) followed by NK maturation markers (CD11b and CD27) to distinguish immature (Imm) and mature (M1 and M2) NK cells, and analysed for CD45 expression. Histograms are representative of n=8 Cd45^fl/fl^Ncr1^iCre/+^ mice and n=7 Cd45^+/+^Ncr1^iCre/+^ mice. **(B-C)** Splenic and liver cells from Cd45^+/+^ (CD45.2; green), Cd45^+/+^ (yellow), Cd45^+/+^Ncr1^iCre/+^ (blue), Cd45^fl/fl^Ncr1^+/+^ (pink), and Cd45^fl/fl^Ncr1^iCre/+^ (purple) mice were analysed for NK cell (CD3^−^NK1.1^+^NKp46^+^) numbers and maturation. (**B)** Splenic NK cell numbers. **(C)** Liver NK cell numbers. **(E)** Frequencies of splenic NK cell maturation populations based on CD11b and KLRG1 expression. **(B-D)** Combined data from 2 independent experiments; each symbol represents an individual mouse. n=4-6 mice. Statistical significance determined by two-way ANOVA with Sidak’s multiple comparisons test.

In contrast to global deletion of *Cd45*, conditional deletion of *Cd45* in NK cells did not result in expansion of the NK cell compartment. NK cell numbers in the spleen and liver of *Cd45^fl/fl^Ncr1^iCre/+^* mice were comparable to *Cd45^fl/fl^Ncr1*^+/+^ and *Cd45^+/+^Ncr1^iCre/+^* control mice, and within a normal range (**Figure 3C & D**). This was not due to an inability to detect NK cells, as a CD45 negative NK population was present in *Cd45^fl/fl^Ncr1^iCre/+^* mice (CD3^−^NK1.1^+^NKp46^+^) throughout NK cell maturation (**Figure 3A & E**). This further suggested that the dramatic increase in NK cell numbers observed in *Cd45^−/−^* mice resulted from changes in early NK development, prior to expression of NKp46.

Interestingly, NK cell maturation does appear to be intrinsically regulated by CD45, as the maturation profile of splenic NK cells from *Cd45^fl/fl^Ncr1^iCre/+^*mice remained skewed to the more mature M2 population (**Figure 3E**), consistent with previous observations (**Figure 1I**).

### CD45 regulates NK cell development

To assess early NK development, NK progenitor populations NKP (CD3e^−^CD8^−^CD4^−^CD19^−^TER119^−^ Ly6G^−^CD11b^−^Ly6D^−^NK1.1^−^; 2B4^+^CD27^+^; cKit^−^CD127^+^; Flt3^−^), pre-NKP (NKP; CD122^−^), and rNKP (NKP; CD122^+^) were enumerated in bone marrow of *Cd45^−/−^* mice (**Figure 4A**). Notably, the absolute number of viable bone marrow cells in *Cd45^−/−^* mice was decreased compared to control mice (**Figure 4B**). In addition, a Lin^−^2B4^med^CD27^med^ population (not previously characterized), was absent in *Cd45^−/−^* bone marrow (**Figure 4A**). The common lymphoid progenitor (CLP; Lin^−^2B4^+^CD27^+^ cKit^−^ CD127^+^ Flt3^+^) population, which gives rise to the pre-NKP and rNKP populations, was marginally increased in bone marrow from *Cd45^−/−^* mice (**Figure 4C**). Lastly, a ∼4 and ∼3-fold decrease in pre-NKP and rNKP populations, respectively, was observed in *Cd45^−/−^* bone marrow (**Figure 4C-D**). This suggested that in *Cd45^−/−^* mice there was a faster transition from CLP through pre-NKP and rNKP, to result in increased numbers of mature M2 NK cells (**Figure 1I**).

**Figure 4.**
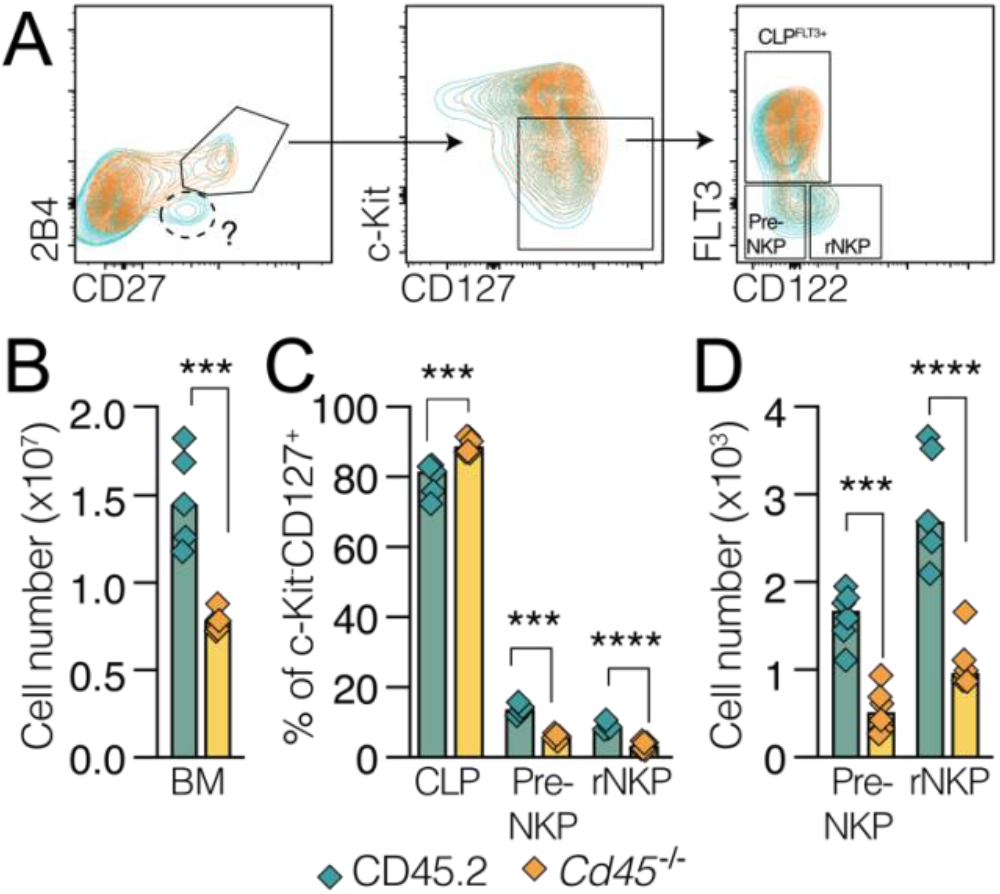
Loss of CD45 reduces NK progenitors in mice. Bone marrow cells were harvested from Cd45^+/+^ (CD45.2; green) and Cd45^−/−^ (yellow) mice and NK cell progenitors analysed by flow cytometry. **(A)** Gating strategy for analysis of NK progenitors post-exclusion of other lineages (CD3e, CD8, CD4, CD19, TER119, Ly6G, CD11b, Ly6D and NK1.1). NK cell populations were analysed based on expression of 2B4, CD27, cKit and CD127. FLT3 expression identifies the common lymphoid progenitor (CLP; CD3e^−^CD8^−^CD4^−^CD19^−^TER119^−^Ly6G^−^CD11b^−^Ly6D^−^NK1.1^−^; 2B4^+^CD27^+^; cKit^−^ CD127^+^; Flt3^+^), while lack of FLT3 expression defines NK progenitors (NKP: CD3e^−^CD8^−^CD4^−^CD19^−^ TER119^−^Ly6G^−^CD11b^−^Ly6D^−^NK1.1^−^; 2B4^+^CD27^+^; cKit^−^CD127^+^; Flt3^−^). Two NK progenitor populations have been defined, the pre-NKP (NKP; CD122^−^) and restricted NKP (rNKP: NKP; CD122 ^+^). An uncharacterized population (Lin^−^2B4^med^CD27^med^) that is missing in Cd45^−/−^ bone marrow is highlighted. **(B)** Enumeration of viable bone marrow (BM) lymphocytes. **(C)** Frequencies of CLP, pre-NKP and rNKP populations. **(D)** Absolute cell numbers. Two independent experiments (n=3 in each experiment). Significance was determined by two-way ANOVA with Sidak’s multiple comparisons test.

Given the increase in CLP cells, the hematopoietic compartment of *Cd45^−/−^* mice was characterized in greater detail. Lineage negative cells (Lin^−^: CD2^−^CD4^−^CD8^−^Gr1^−^F4/80^−^CD19^−^B220^−^Ly6G^−^TER119^−^ NK1.1^−^) were analysed for progenitor and stem cells (**Figure 5A**). Again, despite a decrease in the total number of viable bone marrow cells in *Cd45^−/−^*mice (**Figure 5B**), the Lin^−^ population was increased (**Figure 5C**). CLPs were characterized by two gating methods (CLP: Lin^−^Sca1^med^cKit^med^IL-7R^+^; CLP2: Lin^−^FSC^lo^cKit^med^IL-7R^+^) [53] and enumerated; both methods confirmed an increased number of CLPs in *Cd45^−/−^* mice (**Figure 5D**). No differences in population frequency or enumeration were observed within the myeloid progenitor compartment (Lin^−^Sca1^−^cKit^+^; **Figure 5E**). The Lin^−^ Sca1^+^cKit^+^ (LSK) population, which contains the hematopoietic stem cells, was marginally increased in *Cd45^−/−^* mice (**Figure 5F**). A ∼2-fold increase was observed in the following LSK stem cell populations: long term hematopoietic stem cells (LT-HSC; CD41^−^CD34^−^LSKs), restricted hematopoietic progenitor 1 cells (HPC-1; CD48^+^CD150^−^LSKs), HPC-1 (CD48^+^CD150^−^LSKs) and hematopoietic stem and progenitor cells (HSPC; CD135^−^FSC^mid^; **Figure 5G-I**). Although the differences in the HSC compartment of *Cd45^−/−^* mice appeared modest, they were consistent with CD45 acting to restrict progenitor and stem cell populations.

**Figure 5.**
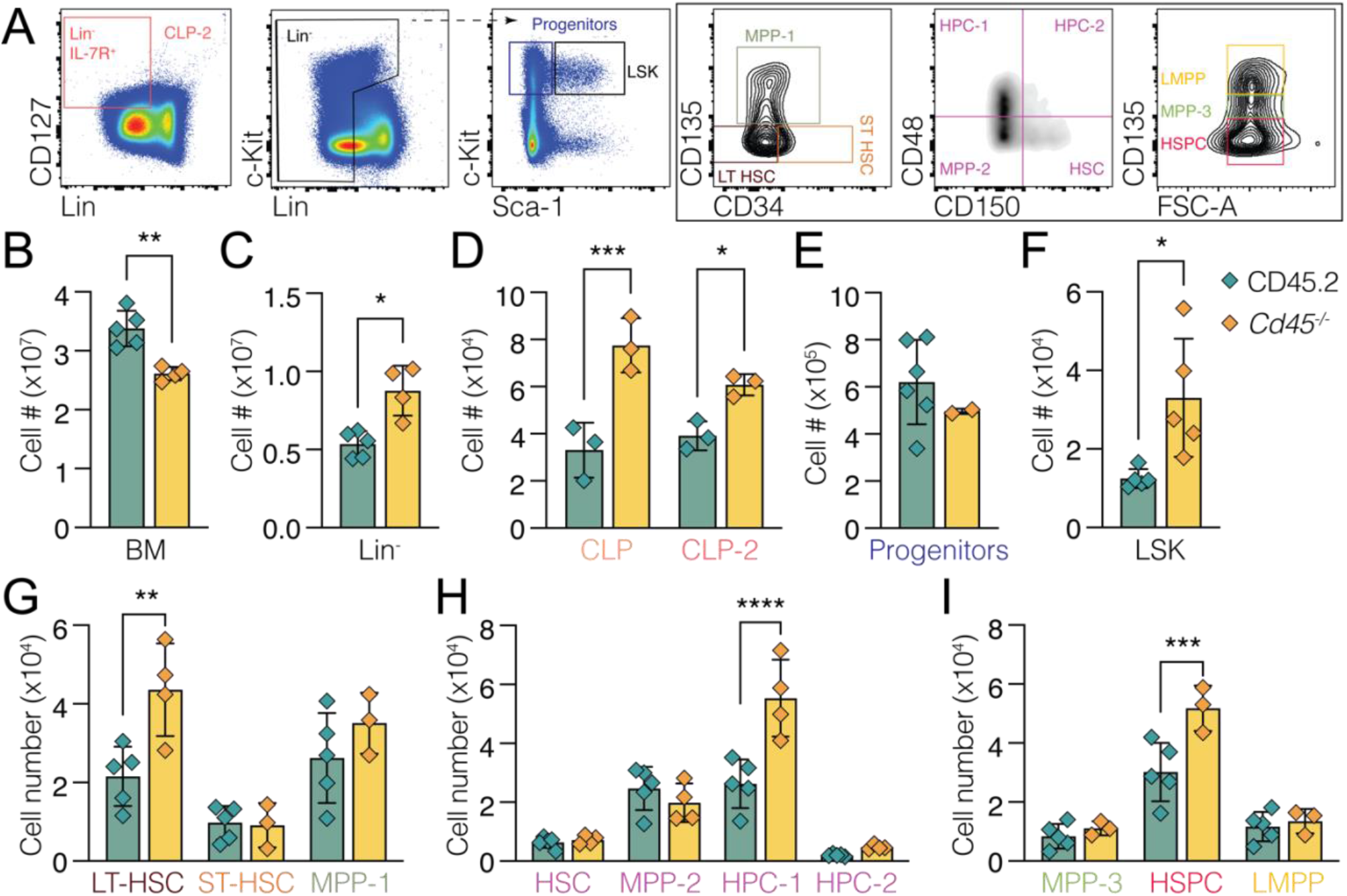
CD45 acts to restrict progenitor and stem cell populations. Bone marrow cells were harvested from Cd45 ^+/+^ (CD45.2; green) and Cd45^−/−^ (yellow) mice for flow cytometric analysis of the HSC compartment. **(A)** Gating strategy to identify populations in the hematopoietic stem cell compartment. Lineage markers (Lin) included CD2, CD4, CD8, Gr1, F4/80, CD19, B220, Ly6G, TER119, and NK1.1. (B-H) Enumeration of **(B)** viable bone marrow cells, (C) viable lineage negative BM cells, (D) the common lymphoid progenitor (CLP) using two recognized gating strategies (CLP: Lin^−^Sca1^med^cKit^med^IL-7R^+^; and CLP2: Lin^−^FSC^lo^cKit^med^IL-7R^+^), (E) Myeloid/erythroid progenitors (Lin^−^ Sca1^−^cKit^+^), (F) the LSK (Lin^−^Sca1^+^cKit^+^) population and (G-I) subsequent hematopoietic stem cell populations determined by flowcytometry and counting beads (123count eBeads). Representative experiment shown of two independent experiments (n=4 mice in each experiment). Significance determined by two-way ANOVA with Sidak’s multiple comparisons test. Abbreviations: Lineage negative cells (Lin^−^: CD2^−^CD4^−^CD8^−^ Gr1^−^F4/80^−^CD19^−^B220^−^Ly6G^−^TER119^−^NK1.1^−^), common lymphoid progenitors (CLP; Lin^−^Sca1^med^cKit^med^IL-7R^+^ and CLP2: Lin^−^FSC^lo^cKit^med^IL-7R^+^), LSK (Lin^−^Sca1^+^cKit^+^), long term hematopoietic stem cells (LT-HSC; CD135^−^CD34^−^LSKs), short term hematopoietic stem cells (ST-HSC; CD135^−^CD34^+^LSKs), multipotent progenitor 1 (MPP-1; CD135^+^CD34^−^LSKs), restricted hematopoietic progenitor 1 cells (HPC-1; CD48^+^CD150^−^LSKs), HPC-2 (CD48^+^CD150^+^LSKs), MPP-2 (CD48^−^CD150^−^LSKs), hematopoietic stem cell (HSC; CD48^−^CD150^+^LSKs), MPP-3 (CD135^mid^FSC^mid^ LSK), lymphoid primed MPPs (LMPP; CD135^hi^FSC^med^LSK), and hematopoietic stem and progenitor cells (HSPC; CD135 ^−^FSC^med^LSK).

Collectively, the data suggest that loss of CD45 likely impacts two stages in early hematopoietic development, resulting in modestly increased LSK/LT-HSC and common lymphoid progenitor populations, and contributing to the expansion of the NK cell compartment in *Cd45^−/−^* mice.

## Discussion

CD45 could be viewed as an immune checkpoint in NK cells, due to the increased NK cell expansion observed in *Cd45^−/−^*mice [1, 2]. Here we provided evidence that the increased NK cell expansion in mice with global loss of CD45, stemmed from a defect in an early progenitor/stem cell population, rather than from a specific effect of CD45 on NK cell proliferation or survival. This is supported by the reduced proliferative capacity of *Cd45^−/−^* NK cells *ex vivo*, the lack of expansion following adoptive transfer of mature *Cd45^−/−^*NK cell populations into lymphoid deficient mice, and finally, conditional deletion of CD45 in NK cells, which had no impact on NK cell homeostasis or the ability of transferred NK cells to expand and persist *in vivo*. In addition, we demonstrated that the NK cell hyperplasia in *Cd45^−/−^* mice was not driven by differential responsiveness to IL-15, a cytokine required for NK cell survival, proliferation, and maturation, nor was different abundance of systemic IL-15 in the in *Cd45^−/−^*mice a likely cause.

Interestingly, although the NK cell expansion observed in *Cd45^−/−^* mice was not present in mice with NK cell specific-deletion of *Cd45*, the skewing of NK cell maturation towards a mature M2 stage was maintained. Again, this was independent of IL-15 levels and suggests that CD45 limits NK cell maturation towards an M2 effector stage, via a yet unknown mechanism. This observation may also account for the apparent reduction in NK cell proliferation observed *ex vivo*. Mature M2 stage NK cells have limited proliferative capacity compared to immature or M1 stage NK cells [1, 54]. Given M2 cells predominate in NK cell pools purified from *Cd45^−/−^* mice, it is perhaps not surprising that we observed significantly reduced *Cd45^−/−^*NK cell proliferation *in vitro*. Again, this is clearly disconnected from the *in vivo* phenotype where *Cd45^−/−^* M2 NK cells are clearly cycling faster than control M2 NK cells as determined by BrdU uptake [1].

The analysis of early progenitor populations revealed a potential role for CD45 in limiting the progenitor/stem pool. Previously, we did not observe a defect in the *Cd45^−/−^* NK precursor population [2], identified as CD122^+^NK1.1^−^DX5^−^CD3^−^CD19^−^ defined by Rosmaraki *et al.*, [55]. In this study, we refined our characterization of the pre-NKP and rNKP populations using markers described by Carotta *et al.,* [56] and Fathman *et al.,*[57] to reveal a defect within the pre and restricted NK progenitor populations [56, 57]. In addition, we observed an increase within both LSK and CLP *Cd45^−/−^*populations, which was not captured by hematopoietic analysis of exon 6-targeted CD45 deficient mice [2]. However, exon 6-targeted CD45 deficient mice retain a small thymic population that expresses CD45, suggesting the lack of perturbation in the lymphoid compartment may be due to residual CD45 expression [4].

## Conclusions

This study attributed the perturbed NK cell homeostasis in CD45-deficient mice to developmental defects impacting the LT-HSC and CLP populations. In CD45-deficient mice, most likely the increase in a common progenitor population, coupled with a faster transit through NK cell development and maturation, manifests as increased numbers of circulating mature NK cells.

Although systemic use of CD45 inhibitors in the clinic would abrogate TCR and BCR signalling [28, 29], both of which are required for an effective anti-cancer immune response, this study identifies a potential clinical application in adoptive immunotherapy. We have shown that CD45 deficiency in the HSC/CLP compartments results in increased NK cell numbers. This predicts that inhibition of CD45 or deletion of the *ptrprc/Cd45* gene, could enhance the yield of NK cells derived from CD34^+^ umbilical cord blood cells or inducible pluripotent stem cells (iPSCs). Improving the yield of NK cells would help address one of the technical limitations currently associated with this approach. Critically, further work is needed to confirm that these observations translate to human NK cell development and do not negatively impact NK cell cytotoxicity.

## Abbreviations

NK: Natural Killer
PTP: protein tyrosine phosphatase
TCR: T cell receptor
BCR: B cell receptor
JAK: Janus kinase
Ig: immunoglobulin
IFN: interferon
PBS: phosphate buffered saline
BSA: bovine serum albumin
I.V.: intravenous
CTV: CellTrace Violet
BM: bone marrow
PI: propidium iodide
CRISPR: Clustered Regularly Interspaced Short Palindromic Repeats
FBS: fetal bovine serum
RT: room temperature
HSC: hematopoietic stem cells
LT-HSC: long term HSC
CLP: common lymphoid progenitors
MPP: multipotent progenitor
LMPP: lymphoid primed MPPs
HPC: hematopoietic progenitor cell
HSPC: hematopoietic stem and progenitor cells

## Acknowledgements

The authors acknowledge the Wurundjeri people of the Kulin nation as the traditional owners and guardians of the land on which most of the work was performed. We thank Prof. Warren Alexander for providing guidance and reagents, Prof. Eric Vivier for providing Ncr1^iCre^ mice and Dr Milon Pang for providing experimental advice. We thank Bioservices staff, Tania Camillari, Natasha Blasch and Sophia Russo, for mouse husbandry and injections, and the Monash Genome Modification Platform for generation of floxed *Cd45* allele mice.

## Author contributions

L.G.M.G designed and executed experiments, and co-wrote the manuscript; CDH designed and performed progenitor analyses; GMB, EL and ZS performed experiments; WG helped design and supervise early experiments; TK contributed experiments that were not included in the final manuscript; JR designed and supervised the study; JEV supervised the study; S.EN supervised the study and co-wrote the manuscript; N.D.H. initiated, designed and supervised the study; All authors reviewed and approved the manuscript.

## Availability of data and materials

All data generated from this study, if not included in this article, are available from the corresponding authors on reasonable request.

## Supplementary data

**Supplementary Figure 1.**
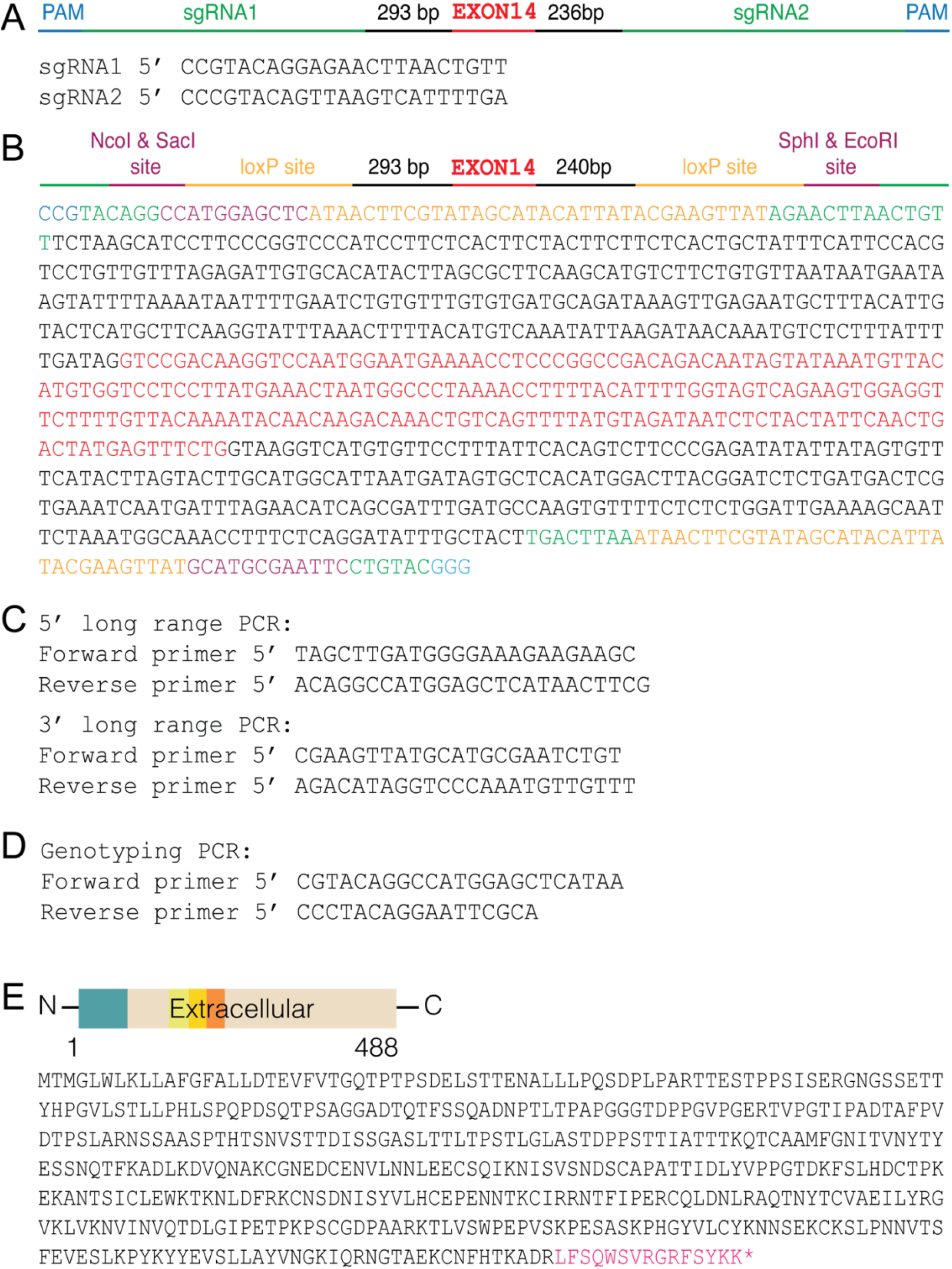
Generation of mice carrying the conditional ptprc allele. **(A)** Schematic color-coded location of sgRNAs relative to exon 14 and guide sequences. **(B)** Schematic representation of donor template used to introduce loxP sites and enzyme restriction sites that flank exon 14 and sequence. **(C)** Primers used to amplify 5’ and 3’ insertion regions to determine s pecificity of donor template integration. **(D)** Genotyping primers used to amplify donor template region and confirm both loxP sites were inserted in one allele. **(E)** Schematic color-coded representation of translated open reading frame post-cre/lox recombination of exon 14 and amino acid sequence

**Supplementary Table 1.**
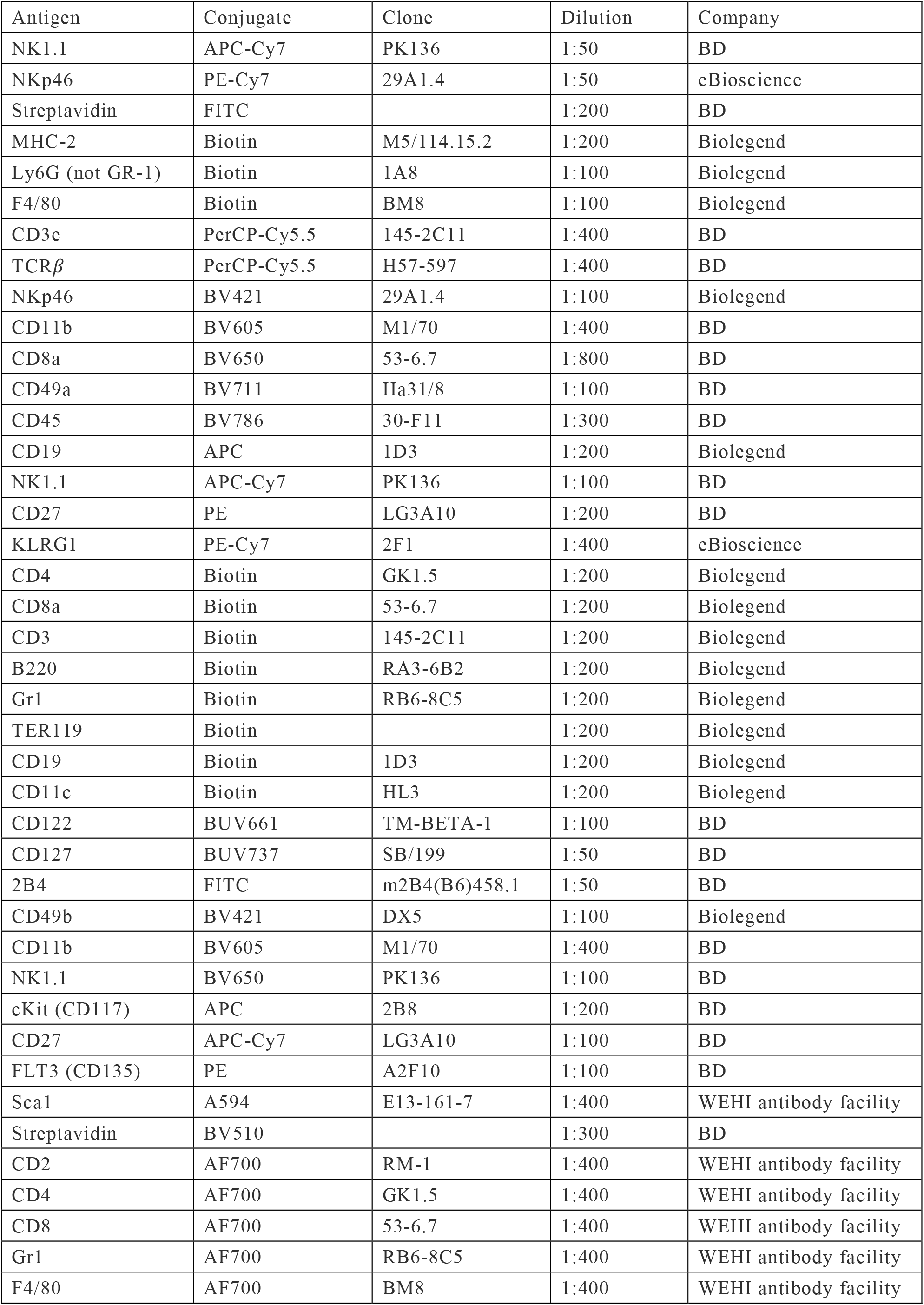
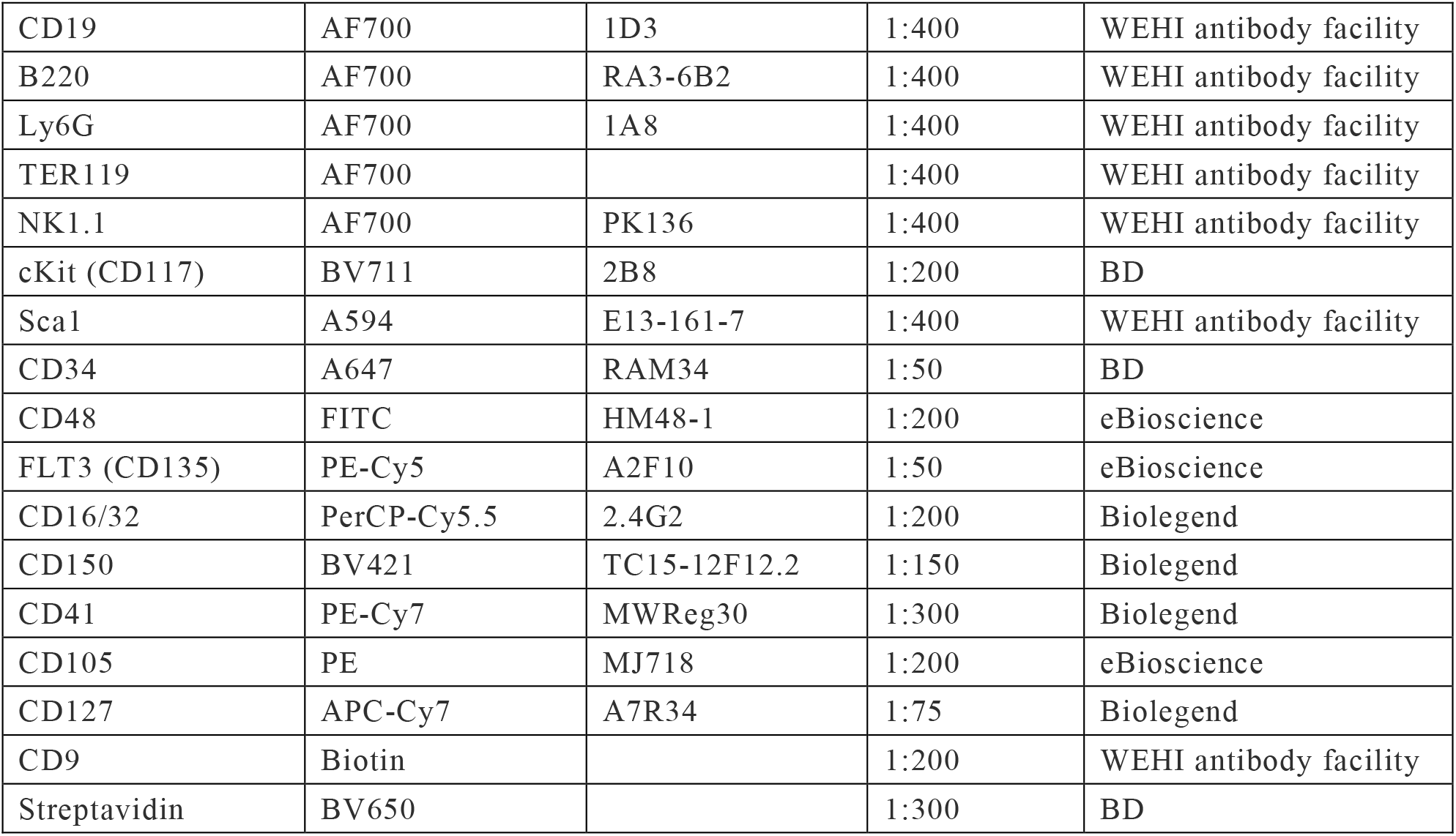
Antibodies used for flow cytometric analysis.

| Antigen | Conjugate | Clone | Dilution | Company |
| --- | --- | --- | --- | --- |
| NK1.1 | APC-Cy7 | PK136 | 1:50 | BD |
| NKp46 | PE-Cy7 | 29A1.4 | 1:50 | eBioscience |
| Streptavidin | FITC |  | 1:200 | BD |
| MHC-2 | Biotin | M5/114.15.2 | 1:200 | Biolegend |
| Ly6G (not GR-1) | Biotin | 1A8 | 1:100 | Biolegend |
| F4/80 | Biotin | BM8 | 1:100 | Biolegend |
| CD3e | PerCP-Cy5.5 | 145-2C11 | 1:400 | BD |
| TCRβ | PerCP-Cy5.5 | H57-597 | 1:400 | BD |
| NKp46 | BV421 | 29A1.4 | 1:100 | Biolegend |
| CD11b | BV605 | M1/70 | 1:400 | BD |
| CD8a | BV650 | 53-6.7 | 1:800 | BD |
| CD49a | BV711 | Ha31/8 | 1:100 | BD |
| CD45 | BV786 | 30-F11 | 1:300 | BD |
| CD19 | APC | 1D3 | 1:200 | Biolegend |
| NK1.1 | APC-Cy7 | PK136 | 1:100 | BD |
| CD27 | PE | LG3A10 | 1:200 | BD |
| KLRG1 | PE-Cy7 | 2F1 | 1:400 | eBioscience |
| CD4 | Biotin | GK1.5 | 1:200 | Biolegend |
| CD8a | Biotin | 53-6.7 | 1:200 | Biolegend |
| CD3 | Biotin | 145-2C11 | 1:200 | Biolegend |
| B220 | Biotin | RA3-6B2 | 1:200 | Biolegend |
| Gr1 | Biotin | RB6-8C5 | 1:200 | Biolegend |
| TER119 | Biotin |  | 1:200 | Biolegend |
| CD19 | Biotin | 1D3 | 1:200 | Biolegend |
| CD11c | Biotin | HL3 | 1:200 | Biolegend |
| CD122 | BUV661 | TM-BETA-1 | 1:100 | BD |
| CD127 | BUV737 | SB/199 | 1:50 | BD |
| 2B4 | FITC | m2B4(B6)458.1 | 1:50 | BD |
| CD49b | BV421 | DX5 | 1:100 | Biolegend |
| CD11b | BV605 | M1/70 | 1:400 | BD |
| NK1.1 | BV650 | PK136 | 1:100 | BD |
| cKit (CD117) | APC | 2B8 | 1:200 | BD |
| CD27 | APC-Cy7 | LG3A10 | 1:100 | BD |
| FLT3 (CD135) | PE | A2F10 | 1:100 | BD |
| Sca1 | A594 | E13-161-7 | 1:400 | WEHI antibody facility |
| Streptavidin | BV510 |  | 1:300 | BD |
| CD2 | AF700 | RM-1 | 1:400 | WEHI antibody facility |
| CD4 | AF700 | GK1.5 | 1:400 | WEHI antibody facility |
| CD8 | AF700 | 53-6.7 | 1:400 | WEHI antibody facility |
| Gr1 | AF700 | RB6-8C5 | 1:400 | WEHI antibody facility |
| F4/80 | AF700 | BM8 | 1:400 | WEHI antibody facility |
| CD19 | AF700 | 1D3 | 1:400 | WEHI antibody facility |
| B220 | AF700 | RA3-6B2 | 1:400 | WEHI antibody facility |
| Ly6G | AF700 | 1A8 | 1:400 | WEHI antibody facility |
| TER119 | AF700 |  | 1:400 | WEHI antibody facility |
| NK1.1 | AF700 | PK136 | 1:400 | WEHI antibody facility |
| cKit (CD117) | BV711 | 2B8 | 1:200 | BD |
| Sca1 | A594 | E13-161-7 | 1:400 | WEHI antibody facility |
| CD34 | A647 | RAM34 | 1:50 | BD |
| CD48 | FITC | HM48-1 | 1:200 | eBioscience |
| FLT3 (CD135) | PE-Cy5 | A2F10 | 1:50 | eBioscience |
| CD16/32 | PerCP-Cy5.5 | 2.4G2 | 1:200 | Biolegend |
| CD150 | BV421 | TC15-12F12.2 | 1:150 | Biolegend |
| CD41 | PE-Cy7 | MWReg30 | 1:300 | Biolegend |
| CD105 | PE | MJ718 | 1:200 | eBioscience |
| CD127 | APC-Cy7 | A7R34 | 1:75 | Biolegend |
| CD9 | Biotin |  | 1:200 | WEHI antibody facility |
| Streptavidin | BV650 |  | 1:300 | BD |

